# Micropollutant-driven bacterial adaptation enables resilient pharmaceuticals biodegradation at trace concentrations in biologically treated wastewater

**DOI:** 10.64898/2026.01.13.699256

**Authors:** Francesca Demaria, Marcel Suleiman, Rafael Bargiela, Manuel Ferrer, Silvia Blázquez Hernández, Abraham Esteve Núñez, Owen L. Petchey, Philippe François-Xavier Corvini, Pilar Junier

## Abstract

Pharmaceutical residues are persistent contaminants that resist conventional wastewater treatment and can disrupt ecosystems; however, microorganisms provide a promising biobased solution to transform or mineralize these complex xenobiotics. Whether pollutant-adapted communities maintain their degradative capacity under realistic environmental conditions remains a long-standing debate in environmental biotechnology. Here, microbial consortia enriched in six membrane bioreactors under high pharmaceutical concentration (100 mg/L) retained full biodegradation capacity across a 5000-fold concentration range. After prolonged exposure to six model compounds (atenolol, caffeine, diclofenac, enalapril, ibuprofen, and paracetamol) complete removal occurred for all except diclofenac. Degradation remained efficient even at lower and environmentally relevant concentrations (1 mg/L-20 µg/L) and recovered rapidly upon re-exposure to higher loads (100 mg/L). Metagenomic profiling revealed enrichment of oxygenase-mediated catabolic pathways supporting this resilience. When transferred to a 7 liters bioreactor treating real wastewater, the adapted community removed targeted and untargeted pharmaceuticals, demonstrating robustness, scalability, and strong potential for sustainable micropollutant remediation.

**Environmental Implication:** Pharmaceuticals and their metabolites are environmentally hazardous because these bioactive micropollutants are persistent and continuously discharged via wastewater, thereby endangering both ecosystem and human health. This study shows that pollutant-adapted microbial consortia can address this challenge, retaining strong degradative function across large concentration fluctuations, including environmentally relevant levels. It also demonstrates scalability: the adapted community can be transferred to real-wastewater operation to remove both targeted and additional pharmaceuticals, supporting a bio-based “polishing” step for wastewater treatment plants. Overall, these findings support more sustainable biological mitigation strategies to reduce micropollutant loads.

## INTRODUCTION

Pharmaceuticals have become ubiquitous micropollutants in aquatic environments, with reported concentrations typically ranging from nanograms to micrograms per liter (Kallenborn et al., 2008; Kümmerer, 2010; Svensson Grape et al., 2023; Tang et al., 2021; Wilkinson et al., 2022) . Their widespread occurrence results from incomplete metabolism in the human body, improper disposal by consumers and manufacturers, and their increasing global consumption (Demaria et al., 2025). Their chemical structures, including ether bonds, benzene rings, and halogenated groups, make pharmaceuticals recalcitrant to biodegradation (Zhang et al., 2008), partially explaining their frequent detection in aquatic ecosystems (Aydin et al., 2019). Moreover, the low environmentally concentrations (µg/L) of pharmaceuticals make them an unfavorable carbon source for microorganisms in wastewater treatment plants (WWTPs), resulting in incomplete biodegradation and their eventual release into the environment (Di Marcantonio et al., 2020; Wang et al., 2021). In wastewater treatment plants, this challenge is further compounded by the strong temporal fluctuations in influent pharmaceutical concentrations, which can vary by several orders of magnitude depending on consumption patterns, hydraulic load, and seasonal effects (Di Marcantonio et al., 2020). However, it is essential to remove these trace concentrations, as pharmaceuticals are biologically active compounds that can exert chronic effects on aquatic organisms and disrupt ecosystem functioning even at very low concentration levels. The accumulation of pharmaceuticals in the environment can have irreversible negative impacts for both ecosystems and human health (Gorovits et al., 2020; Persson et al., 2013; Steffen et al., 2015). Pharmaceuticals detected in the environment span various categories. Caffeine, a central nervous system stimulant, is common in sewage due to its high consumption (Khasawneh and Palaniandy, 2021; Wu et al., 2012; Zhou et al., 2016); analgesics and anti-inflammatory pharmaceuticals such as ibuprofen, paracetamol and diclofenac are also widespread and often sold without prescription (dos S. Grignet et al., 2022; Zhou et al., 2016). Atenolol, a β-blocker, and enalapril, used for heart failure are commonly detected in treated wastewater (Lavén et al., 2009; Ruan et al., 2019; Salgado et al., 2010).

Bio-based solutions are promising complementary techniques that can improve the removal of pharmaceuticals from wastewater (Demaria et al., 2025). In a recent study, a synthetic biology approach was successfully developed to engineer *Vibrio natriegens* to degrade various organic pollutants in industrial wastewater(Su et al., 2025). Here, we propose a complementary top-down strategy based on our previous work, in which we enriched microbial communities using membrane bioreactors (MBRs) to remove high concentrations of the aforementioned pharmaceuticals (Suleiman et al., 2023). This suggested new directions for research; as there is a fundamental difference between the conditions that promote the growth of pollutant-degrading microorganisms (high pollutant concentrations) and the concentrations measured in real wastewater systems (low pollutant concentrations) (Fenner et al., 2021; Yin et al., 2024). Enriching effective degraders typically requires strong and sustained selective pressure, as only under high pollutant loads do microbial communities redirect their metabolic potential toward utilizing pharmaceuticals as growth substrates or co-metabolic targets [23]. ^24^At low concentrations, these compounds provide insufficient energetic or nutritional benefits to sustain such activity, limiting the natural development of specialized degradative pathways (Yin et al., 2024). Consequently, it remains uncertain whether microbial consortia selected under high pollutant stress can retain their biodegradation functionality when confronted with trace and fluctuating environmental relevant concentration. Therefore, we set up a series of experiments mimicking the fluctuating pharmaceutical concentrations typically observed in wastewater treatment plants. In parallel, we aimed to determine whether microbial communities enriched under high pollutant loads could retain their degradative activity when exposed to environmentally relevant low concentrations of these compounds. Finally, we evaluated the performance of the lab-adapted microbial communities in removing a wide range of residual pharmaceuticals from real biologically treated wastewater using a 7 L membrane bioreactor.

## MATERIALS AND METHOD

### Set-up of a Lab-scale Membrane-Bioreactors (MBRs) and experimental design

Six lab-scale MBRs with a total volume of 1 L were filled with 500 mL synthetic wastewater (Suleiman et al., 2023) (Supplementary Materials S1). The MBRs were constantly aerated and stirred. MBRs were equipped with polyethersulfone ultrafiltration membranes (NADIR UP150, Mann+ Hummel) (Fig. 1a, 1b) with a pore size of 0.08 µm with a total membrane area of 30 cm^2^. The MBRs were inoculated with 1% v/v of activated sludge from an in-house MBR treating wastewater from the university campus. The feed was constantly pumped into the vessel with a flow rate of 8 mL/h, resulting in a Hydraulic Retention Time (HRT) (Wu et al., 2022) of approximately 62 hours. The concentration of pharmaceuticals was set as follows: 100 mg/L for 10 weeks, 1 mg/L for 4 weeks, 100 mg/L for 5 weeks (Fig. 1c). The measured Chemical Oxygen demand (COD) concentrations were 171 mg/L (synthetic wastewater), 185 mg/L (synthetic wastewater + 1 mg/L of each pollutant), and 1074 mg/L (synthetic wastewater + 100 mg/L of each pollutant).

**Figure 1.**
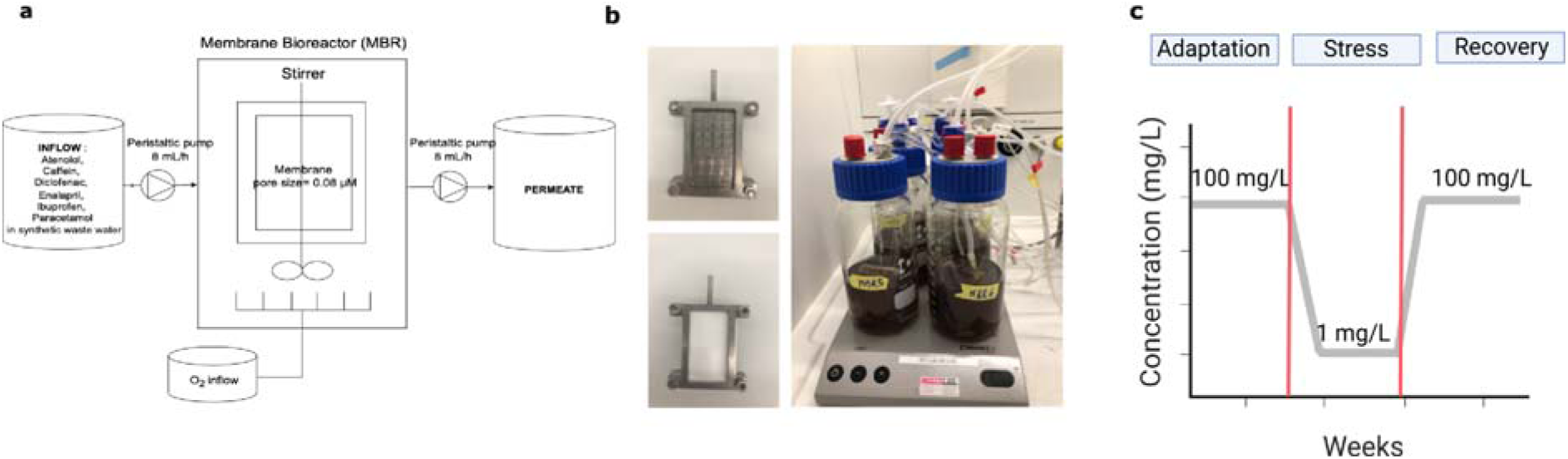
Description of the MBRs structure and experimental design. **a,** Schematic of the MBR system fed with synthetic wastewater containing six pharmaceuticals (atenolol, caffeine, diclofenac, enalapril, ibuprofen, paracetamol); flow rate: 8 mL/h; aerated continuously. **b,** Pictures of the six lab-scale MBRs equipped with 0.08 µm ultrafiltration membranes. **c,** Overview of pharmaceutical concentrations across three phases: adaptation (100 mg/L, 10 weeks), stress (1 mg/L, 4 weeks), and recovery (100 mg/L, 5 weeks).

### Sampling and HPLC analysis

Weekly, 2 mL samples were collected from the MBRs and centrifuged at 17,000 rpm for 10 min. The supernatant was used for high performance liquid chromatography (HPLC) analysis, and the pellet for DNA extraction and sequencing. Pharmaceuticals were quantified by HPLC using a Zorbax SB-C18 column (3.5 µm, 3.0 × 150 mm) with a Zorbax guard column (5 µm, 4.6 × 12.5 mm). The mobile phase consisted of solvent A (water + 0.1% formic acid) and solvent B (methanol), with the following gradient: 0 min A 80%, B 20%; 2 min A 80%, B 20%; 8 min A 20%, B 80%; 12 min A 5%, B 95%; 18 min A 12%, B 95%. Retention times were 1.5 min (atenolol), 2 min (paracetamol), 7 min (caffeine), 10.5 min (enalapril), 13.5 min (diclofenac), and 13.7 min (ibuprofen). Detection was at 230 nm, except 205 nm for enalapril. The removal efficiency for targeted pharmaceuticals and MBR at each sampling point was calculated as following (1):

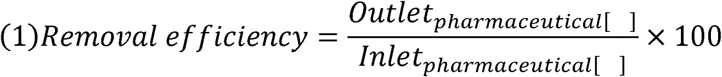

### DNA extraction, sequencing, and data processing

DNA extraction was performed using the ZymoBIOMICS DNA Miniprep Kit (ZymoResearch) according to the manufacturer’s instructions. The V4 region of the 16S rRNA gene was amplified, and DNA libraries were prepared using the Quick-16S™ Plus NGS Library Prep Kit (V4) (ZymoResearch). The 4 pM DNA libraries, each spiked with 25% PhiX, were sequenced in-house using Illumina MiSeq platform following the manufacturer’s protocol. Sequencing data were processed using the dada2 R package (Callahan et al., 2016), with filtering based on primer sequences, quality scores, error rates, and chimera removal. The resulting sequence table was aligned to the SILVA ribosomal RNA database (Quast et al., 2013), version 138 (99% non-redundant dataset). A phyloseq object was generated using the phyloseq R package (McMurdie and Holmes, 2013), incorporating the amplicon sequence variant (ASV) table, taxonomy table, and sample metadata. Further analyses were conducted using the phyloseq (McMurdie Paul J. AND Holmes, 2013) and vegan (Oksanen et al., 2013) R packages. The phyloseq object, metadata, and detailed R code for analysis are available on GitHub (https://github.com/Fra170197) and raw sequencing data can be accessed via NCBI SRA under project IDs: PRJNA1301277 (MBR1) and PRJNA1301261 (MBRs2-6).

### Community analysis

We used ggplot in R studio environment for data visualization. The microbial composition was represented as the relative abundance of microorganisms at genus level in each sample. A set threshold was used to remove all the genus whose abundance was less than 5% of the total. A non-metric multidimensional scaling (NMDS) analysis was used to measure the distance or dissimilarity matrix of each sample in the adaptation phase, and then in the combined adaptation, stress, and recovery phases. The samples distances were calculated using Bray-Curtis distance, the dimensions were reduced at 2 (k=2) and the iterations were 20 taking the smallest stress value, below 0.2. The analysis was performed using the package vegan (Oksanen et al., 2013). The alpha diversity was quantified using Shannon entropy as the default setting of vegan package (Oksanen et al., 2013). Furthermore, the data set was divided in two subgroups to detect the key genera involved in the degradation of pharmaceuticals. We divided the data in **“H”** (high removal efficiency samples) representing all the samples (in adaptation and recovery phase) across the MBRs and the time, in which the degradation of atenolol, enalapril was above 99%; and **“L”** (low removal efficiency samples), representing all the samples in which the degradation of atenolol, enalapril was less than 99%. The 99% threshold was set arbitrarily. Caffeine, paracetamol, and ibuprofen were excluded from the analysis since they were easier to remove at the recovery phase. Once these two groups were defined, the alpha diversity among the two was analyzed using the vegan package (Oksanen et al., 2013). Moreover, the differential abundance analysis was conducted using Deseq2 package (Love et al., 2014), applying a Wald test with a significance threshold of p < 0.01 to identify statistically relevant microbial differences between groups.

### Metagenomics analysis

For metagenomic analysis, the samples at day 36 (acclimatization phase), 104 (stress phase), 136 (recovery phase) from MBR1 (lowest performance on the recovery phase) and MBR6 (high performance) were selected. DNA was extracted using the ZymoBIOMICS DNA Miniprep Kit (ZymoResearch) according to the manufacturer’s instructions. Zymo prepared the sample library using the Illumina DNA Library Prep Kit (Steemers and Gunderson, 2005). Finally, we performed shotgun metagenomic sequencing of the samples on the NovaSeq® platform. The raw sequencing data can be accessed via NCBI SRA under project IDs: PRJNA1362991. Metagenomes were assembled using bbtools 38.90 (Bushnell Brian AND Rood, 2017) for re-pairing reads, artifacts and adapters removal, correction and merging of reads. The final set of reads were assembled using Megahit v1.1.3 (Li et al., 2015). Once the assembly was produced, gene prediction and annotation were performed using Prokka 1.4.5 (Seemann, 2014) (Supplementary Table 1). Metagenome-assembled genomes (MAGs) were calculated for MBR1 and MBR6 metagenomes. Briefly, bins were initially obtained using three different bioinformatics tools: Maxbin2 (Wu et al., 2016), Metabat2 (Kang et al., 2019), and Concoct (Alneberg et al., 2014). Then, MAGs were dereplicated using Dash Tool (Sieber et al., 2018) and purified using MAGsPurify2 (Camargo et al., 2023). Taxonomic classification of MAGs was performed using GTDBtk v2.4.0. and GTDB database v214 (Parks et al., 2022), https://gtdb.ecogenomic.org/) (Supplementary Table 2). Quality control was checked using CheckM2 (Chklovski et al., 2023). This resulted in 376 dereplicated MAGs, out ow which where 108 are considered as being of acceptable quality.

### Adapted Microbial Community Performance at Relevant Pharmaceutical Concentrations

A batch experiment was conducted in triplicate using synthetic wastewater containing 20 µg/L of the targeted pharmaceuticals to evaluate the performance of the adapted microbial community at environmentally relevant pharmaceutical concentrations. The inoculum taken from MBR6 was washed and added to the medium. A control group was included to assess sorption of pharmaceuticals to the biomass during the MBR cultivation phase, by adding the biomass to a medium without pharmaceuticals. Samples were collected immediately after inoculation and on day 3, and pharmaceutical concentrations were analyzed by LC-MS (liquid chromatography-mass spectrometry) (Supplementary Materials S2).

### Set-up of 7 L MBR

A 7-L membrane bioreactor (MBR) (Fig. 2) equipped with a polyethersulfone ultrafiltration membranes with a pore size of 0.08 µm with a total membrane area of 1200 cm^2^, was set up as previously described in *Set-up of Lab-scale Membrane Bioreactors and Experimental Design*. The MBR was operated under aerobic conditions with a working volume of 3.5 L and an HRT of 40 hours. An inoculum of 50 mL from frozen MBR6 was thawed and added to the reactor along with synthetic wastewater containing a 100 mg/L mixture of pharmaceuticals. The reactor was operated in batch for one week. Subsequently, the MBR was operated on a continuous fashion, and the influent (COD_avg_= 250 mg/L) was wastewater biologically pre-treated using a METland® at the IMEDEA facilities, Spain. Influent and effluent samples were collected for subsequent mass spectrometry (MS) analysis.

**Figure 2.**
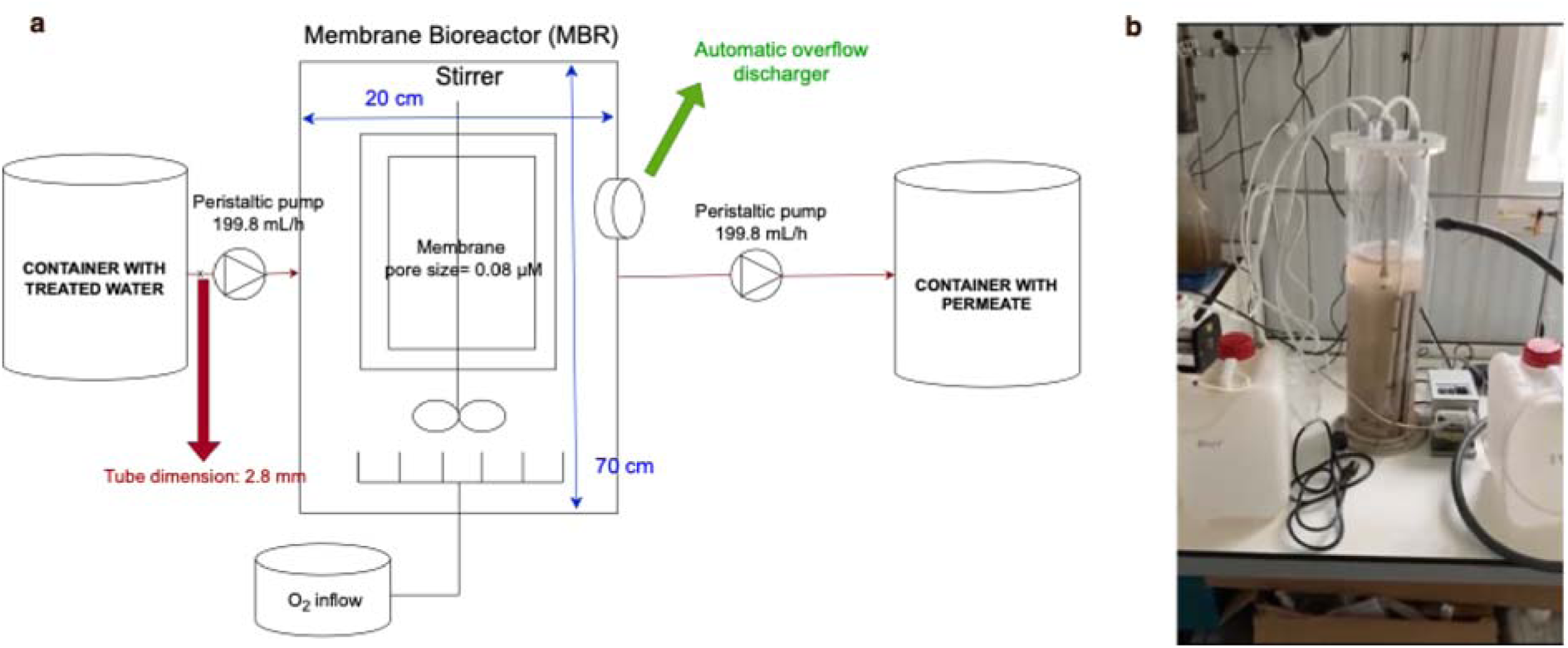
7-liter MBR. **a**, Schematic parameters overview of the scaled-up MBR. **b,** 7-L MBR in continuous operation after inoculation.

### MS-method for 7-L MBR

Two liters of samples (input and output) were taken twice per week and filtered using 0.7 µm glass fiber. Furthermore, they were subjected to a solid phase extraction process applying a preconcentration factor of 100 and were diluted according to instrumental requirements. Pharmaceutical quantification was performed using an Agilent 1200 series high-performance liquid chromatography system coupled to an Agilent 6495 triple quadrupole mass spectrometer equipped with an electrospray ionization (ESI) interface incorporating Jet Stream and i-Funnel technologies. Analyses were conducted in both positive (ESI+) and negative (ESI-) ionization modes to ensure optimal detection of all compounds. For positive ionization, samples were separated on a Kinetex Biphenyl column (50 mm x 3 mm, 2.7 µm) protected by a security guard cartridge. The mobile phase consisted of 0.1% formic acid in ultrapure water (A) and 0.1% formic acid in methanol (B), with a gradient increasing from 2% to 100% B over 30 minutes, maintained for 5 minutes, and followed by a 4-minute re-equilibration. For negative ionization, separation was achieved using a Poroshell 120 EC-C18 column (50 mm x 3 mm, 2.7 µm) with a guard cartridge, employing 1 mM ammonium fluoride in ultrapure water (A) and a methanol–acetonitrile mixture (65:35, v/v) (B). The gradient ramped from 5% to 100% B within 12 minutes, held for 5 minutes, and re-equilibrated for 4 minutes. In both cases, the flow rate was maintained at 0.6 mL/min, the column temperature at 40 °C, and the injection volume at 20 µL. The electrospray source parameters were set as follows: drying gas temperature 250 °C and flow 13 L/min, sheath gas temperature 350 °C and flow 11 L/ min, nebulizer pressure 45 psi, and capillary voltage of +4000 V for ESI⁺ and −3000 V for ESI⁻. Detection was carried out using dynamic multiple reaction monitoring (MRM), with the method divided into 2-minute segments to optimize the number of transitions monitored without compromising sensitivity. Based on the most characteristic and sensitive MS/MS fragments, a linear range of the 30 compounds analyzed (Supplementary Table 1) from 10 ng/L– 40 µg/L was achieved.

## RESULTS

### Adaptation of the lab-scale MBR communities to high-load pharmaceuticals

Six lab-scale MBRs were inoculated with biomass of a MBR system that was treating the selected micropollutants for two months (Suleiman et al., 2023). The reactors were operated in parallel to evaluate the removal of atenolol, caffeine, diclofenac, enalapril, ibuprofen, and paracetamol at influent concentrations of 100 mg/L each (Fig. 3a). Caffeine and paracetamol were completely removed within 7 and 16 days, respectively. Following approximately 29 days of operation, ibuprofen removal efficiency reached 100% in all reactors. A transient loss of activity was observed in MBR1 at day 36, but removal recovered by day 73. In contrast, atenolol and enalapril were more recalcitrant to biodegradation, as their complete removal was achieved only after a prolonged adaptation period of 73 days. Diclofenac was not removed under the tested conditions. To evaluate whether differences in removal efficiency dynamics were associated with microbial composition, we analyzed community profiles over time (Fig. 3b). The NMDS (Non-Metric Dimensional Scaling) analysis (Fig. 3c) reflected distinct adaptation trajectories for each microbial community. Initially, the six microbial communities were dominated by *Achromobacter, Burkholderia,* and *Pseudomonas*, and the communities diversified as adaptation progressed. Specific genera such as *Reyranella* and *Leucobacter* became enriched in all MBRs, except for MBR1. Over time, the community structures of MBR2, MBR3, and MBR6 converged, diverging from those of MBR1 and MBR4, which formed a separate cluster in the NMDS analysis (Fig. 3c). Although diversity initially decreased, its subsequent rise across all reactors, evidenced by the increasing Shannon index (Fig. 3d), suggested that enhanced biodiversity was a key driver of microbial adaptation and pharmaceuticals biodegradation efficiency.

**Figure 3.**
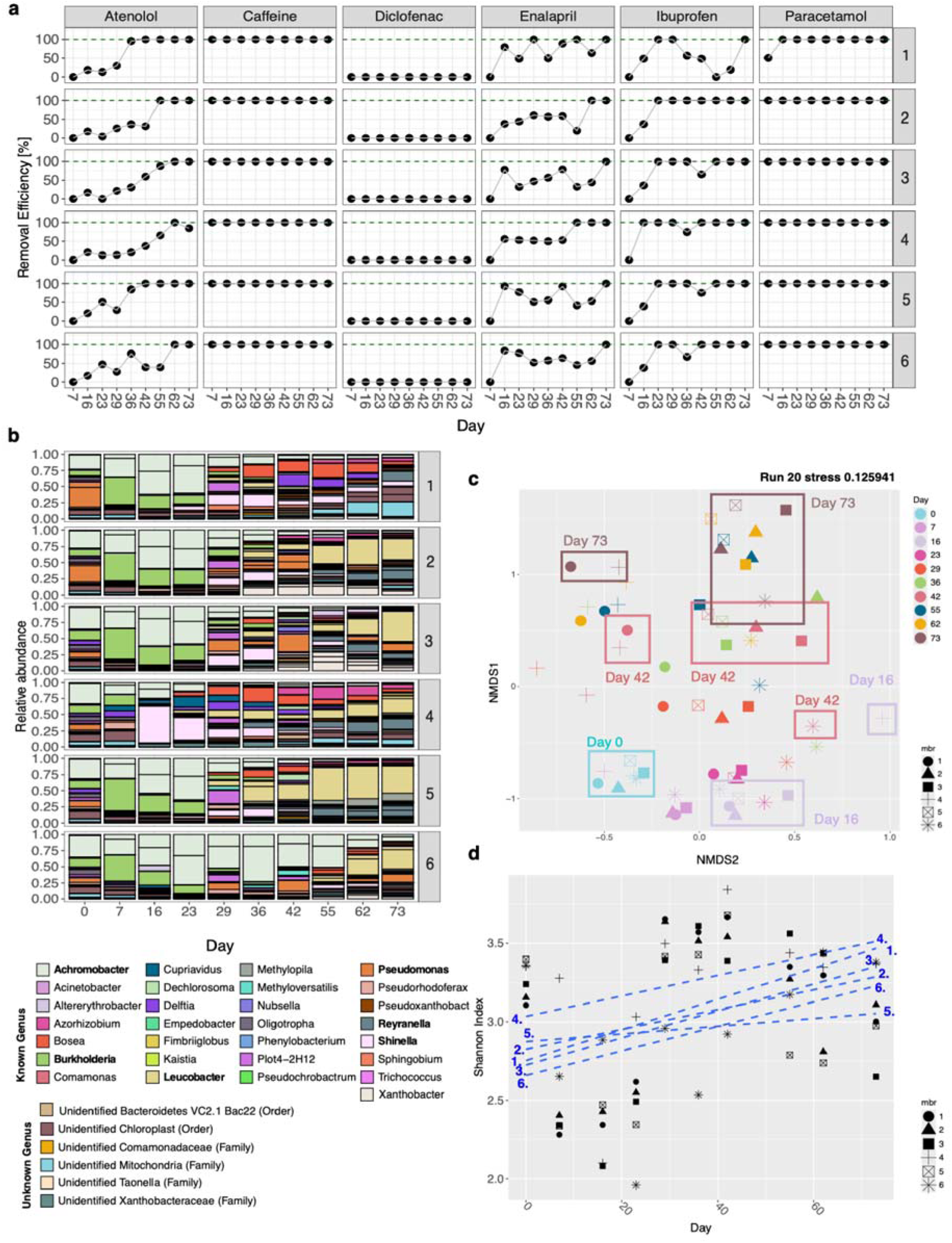
Analysis of the high-load pharmaceutical adaptation phase: a, Removal efficiency of 100 mg/L of atenolol, caffeine, diclofenac, enalapril, ibuprofen, and paracetamol over 73 days in six MBRs. b, Relative abundance of microbial genera that are above 5% across all samples over the operation time; genera in bold were the most relevant. c, NMDS showing community development in six lab-scale MBRs (different shapes) from day 0 and 73; stress = 0.13 (20 iterations). D, Shannon diversity of microbial communities in the six MBRs from day 0–73.

### Lab-scale MBR communities resiliency to stress

After 73 days of incubation with high-load pharmaceuticals, the concentration was switched to 1 mg/L in all the MBRs, which is defined as “stress phase”. During the stress phase, removal efficiency of caffeine, enalapril, ibuprofen, and paracetamol remained stable at 100% in all MBRs for 30 days. Only atenolol showed a temporary decrease in removal efficiency (day 104, MBR1, 2, 4,and 6). In the recovery phase (when to pharmaceutical concentrations were restored to 100 mg/L for 30 days), caffeine, ibuprofen, and paracetamol were fully removed. However, atenolol and enalapril exhibited some fluctuations in removal efficiency, particularly in MBR1. Diclofenac persisted throughout, indicating limited degradability of this compound by the microbial communities (Fig. 4a).

**Figure 4.**
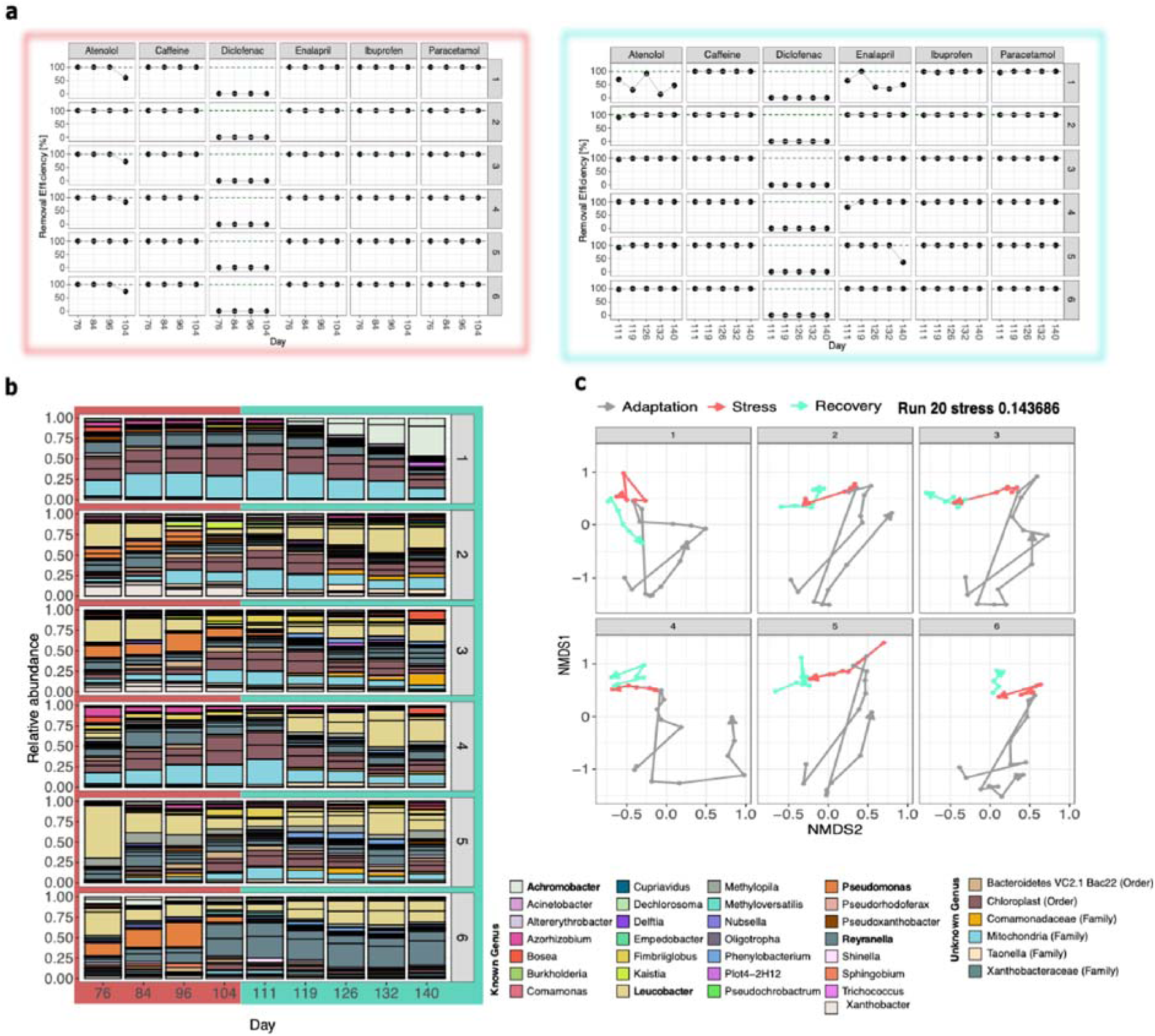
Stress and recovery phases: a, Removal efficiency of six pharmaceuticals in the six lab-scale MBRs during stress (1 mg/L, red frame) and recovery (100 mg/L, turquoise frame). b, Relative abundance of microbial genera that are above 5% during stress and recovery phases; bold indicates taxa more relevant. c, NMDS plot (stress = 0.14) showing community shifts across phases: grey (adaptation), red (stress), turquoise (recovery).

The application of fluctuating pharmaceutical concentrations (100 mg/L-1 mg/L-100 mg/L) significantly altered the microbial community composition. In MBRs with high removal efficiency of pharmaceuticals (MBR 2-6), *Leucobacter* decreased at low pharmaceutical concentrations and increased when concentrations were restored to high concentrations. *Reyranella* abundance increased in MBR6, while *Pseudomonas* declined and disappeared during recovery. The underperforming MBR1 had a distinct community dominated by *Achromobacter* during recovery, along with reduced Shannon diversity (Supplementary Materials S3), highlighting the link between high biodiversity and stable pollutant removal (Fig. 4b).

NMDS analysis illustrated the temporal trajectories of microbial community composition across all MBRs (Fig. 4c). During the adaptation phase, no consistent pattern was observed, as each community evolved independently. However, during the stress and recovery phases, five of the six MBRs (2–6) exhibited similar trajectories, suggesting convergent community responses. In contrast, MBR1 followed a distinct trajectory, likely explaining its lower removal efficiency for atenolol and enalapril during recovery phase. This highlights the importance of microbial community evolution in maintaining functional resilience under fluctuating conditions.

### Genes and genera involved in pharmaceuticals degradation at lab scale level

Shotgun metagenomic sequencing was performed to identify putative genes associated with the biodegradation of atenolol, caffeine, ibuprofen, paracetamol, enalapril, and diclofenac in two membrane bioreactors: MBR1 (low-performance) and MBR6 (high-performance). A total of 181 sequences encoding potential pharmaceutical-degrading enzymes were identified (72 in MBR1, 109 in MBR6) based on homology with experimentally validated enzymes (Supplementary Tables S2-S3). Sequence identity ranged from 60-100%, and hits were found for four of the six target compounds (atenolol, caffeine, ibuprofen, and paracetamol), but none for enalapril and diclofenac. Of these, 157 sequences were assigned to 108 high-quality Metagenome-Asssembled Genome (MAGs), linking functional potential to specific bacterial taxa (Fig. 5a).

**Figure 5.**
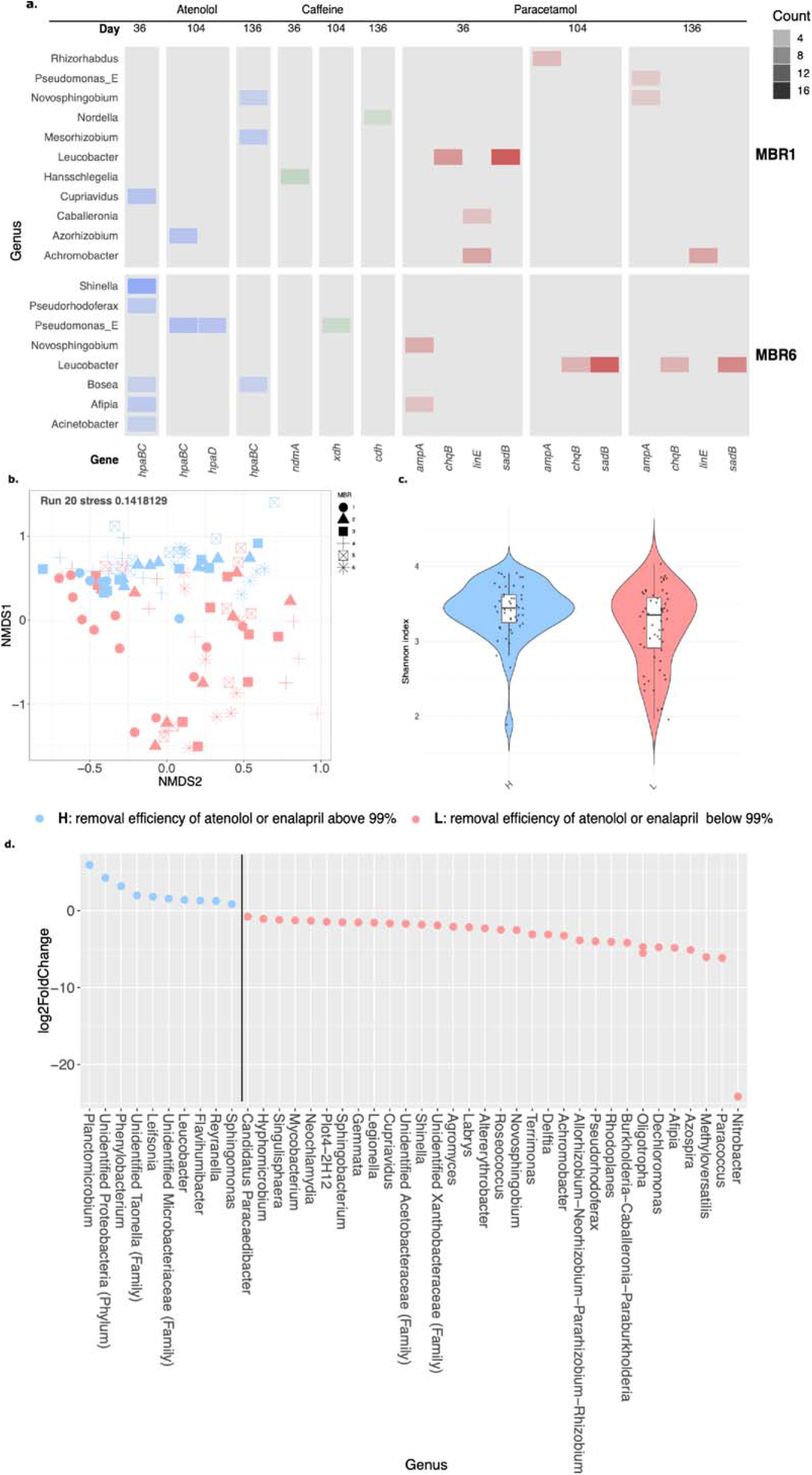
Analysis of key players. **a**, Genes associated to atenolol (blue), caffeine (green), and paracetamol (red) degradation in MBR1 and MBR6 over time, identified via metagenomics. The y-axis shows genera encoding degradation enzymes (MAGs); color shades indicate metagenomic signal intensity based on MAGs hits. **b,** NMDS of two sample groups made across adaptation and recovery: H (blue, >99% removal of atenolol/enalapril) and L (red, <99%). **(c.)** Violin plots of Shannon diversity across H and L groups; boxplots show medians and Interquartile Ranges. Diversity differed significantly (Wilcoxon, *p* < 0.01). **d,** DESeq analysis of taxa abundance between H and L; y-axis lists significantly different genera, x-axis shows log fold change, with symbols marking MBRs.

Genes encoding atenolol-degrading enzymes (*hpaBC*, *hpaD*) were detected in both reactors, with higher relative abundance in MBR6 (day 36, hits > 7.5) (Supplementary Material S4) and mapped to *Shinella*, *Bosea*, and *Acinetobacter.* Paracetamol-related genes (*sadB*, *chqB*, *linE*) were enriched in MBR1, and MBR6 (hits > 10, day 36) and associated with *Leucobacter* and *Achromobacter*. Caffeine-degrading genes (*ndmA–C*, *cdh*) were detected in both MBRs and assigned to *Nordella*, *Hansschlegelia*, and *Pseudomonas*. Genes linked to ibuprofen metabolism (*ipfF*, *ipfA*) were detected at low abundance (∼2.5 hits) in both MBRs (Supplementary Material S4) and could not be mapped to MAGs, suggesting degradation through alternative or low-abundance pathways. Diclofenac-degrading genes (*P450 monooxygenases*) were not detected, consistent with it not being removed.

Atenolol and enalapril exhibited the lowest biodegradability among the tested compounds. To identify taxa potentially involved in their removal, samples were grouped into high-efficiency (H) and low-efficiency (L) communities using a 99% removal threshold for both, atenolol and enalapril (Supplementary Materials S5). NMDS (stress = 0.14) analysis showed that H communities clustered closely, while L communities were more dispersed (Fig. 5b). Higher Shannon diversity (p < 0.01) was significantly associated with high removal efficiency (Fig. 5c). Differential abundance analysis (DESeq2) revealed that *Reyranella* and *Leucobacter* were enriched in efficient communities, whereas *Achromobacter* and *Burkholderia* dominated low-efficiency samples (Fig. 5d).

These results suggest that biodegradation performance is driven by microbial community structure and emphasize the need for deeper functional and pathway-level characterization to link genetic potential with observed degradation capacity.

### Adapted microbial community removal efficiency under environmentally relevant conditions

The biodegradation capacity of the adapted microbial community from lab-scale MBR6 was evaluated under environmentally relevant conditions using synthetic wastewater containing pharmaceuticals with a concentration of 20 µg/L (Supplementary Material S6). The community effectively removed the five target compounds (atenolol, caffeine, enalapril, ibuprofen, and paracetamol) within three days, except for diclofenac, which remained recalcitrant over this period.

To assess its bioremediation potential in relevant environment, the adapted community from MBR6 was inoculated into a 7 L MBR to clean biologically treated wastewater containing residual pharmaceuticals at concentrations ranging from ng/L to µg/L. The system achieved near-complete removal of caffeine, enalapril, ibuprofen, and paracetamol, while atenolol removal fluctuated throughout the month, reaching 75% by day 32. Notably, diclofenac, previously not degraded under laboratory conditions, exhibited up to 50% degradation after 32 days (Fig. 6a).

**Figure 6.**
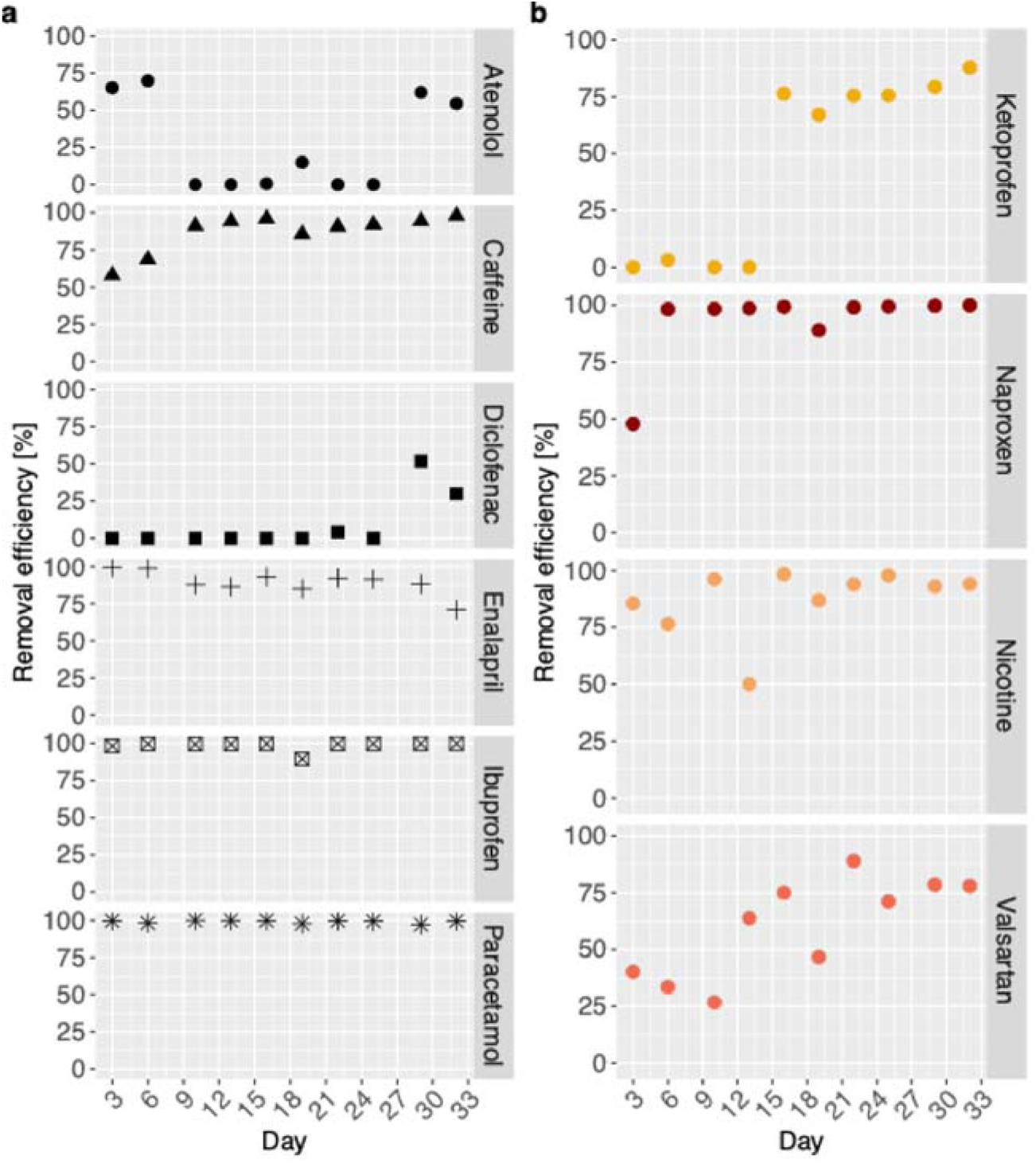
Removal of pharmaceuticals from treated wastewater in a 7-L MBR. The y-axis represents removal efficiency (calculated as described in the materials and method section), while the x-axis indicates operational days. **a,** different shapes denote targeted compounds (right facet), and **b,** different colors represent the untargeted compounds (right facet).

The treated effluent contained additional, previously untargeted pharmaceuticals not present in the initial synthetic feed, including naproxen, ketoprofen, valsartan, and nicotine (Supplementary Table 1). Despite this compositional complexity, the adapted community demonstrated functional flexibility, achieving 75–100% removal of these compounds within 32 days (Fig. 6b).

These findings demonstrate that the adapted microbial community remained functionally robust under realistic operating conditions, underscoring its promise for advanced treatment stages aimed at reducing pharmaceutical residues in treated effluents.

## Discussion

The pervasive occurrence of pharmaceuticals in aquatic environments underscores the urgent need for sustainable and biologically based remediation strategies. Ensuring that adapted consortia remain effective under fluctuating wastewater composition typical of real wastewater systems a key challenge in environmental biotechnology (Svendsen et al., 2020; van Bergen et al., 2021). Operating six membrane bioreactors (MBRs) under fluctuating pharmaceutical concentrations demonstrated that adapted microbial communities could effectively function as an *in situ* biological “polishing step” for wastewater quality improvement. Importantly, the microbial communities enriched under high pollutant pressure retained full biodegradation capacity despite a 5000-fold reduction in substrate concentration (from 100 mg/L to 20 µg/L), indicating strong functional resilience and concentration adaptability. Furthermore, upon re-exposure to high pollutant concentrations (from 1 mg/L to 100 mg/L), the lab-scale MBR communities fully recovered their degradation efficiency, highlighting their resilience and ecological stability in response to fluctuating environmental conditions (Fig. 4). This robustness is critical for real wastewater treatment operations, where influent loads vary widely (Di Marcantonio et al., 2020). Therefore, this work provides experimental evidence that microbial communities enriched under high pollutant concentrations (mg/L range) retain their biodegradation capacity when exposed to environmentally realistic, low-concentration conditions (µg /L range).

All six lab-scale MBRs achieved complete degradation of the five target pharmaceuticals (except diclofenac), and the six microbial communities evolved along distinct compositional trajectories during the adaptation phase (Fig. 3). This indicates that different microbial assemblies can achieve comparable removal efficiencies, highlighting the diverse adaptability of wastewater-associated microbiomes to environmental pressures. However, the improved removal of the more recalcitrant compounds, atenolol and enalapril, was associated with a shift toward more diverse and specialized communities (Fig. 3). DESeq2 analysis revealed that *Leucobacter* and *Reyranella* were significantly more abundant in high-efficiency samples (>99% removal). *Leucobacter* has been previously linked to the hydroxylation of sulfamethoxazole (Reis et al., 2018) and was also detected in MBRs exposed to high pharmaceuticals load (Suleiman et al., 2023). *Reyranella* has been positively correlated with emerging pollutant removal in subsurface wastewater infiltration systems (Peng et al., 2025), highlighting its potential role in the biodegradation of pharmaceuticals. The early dominance of *Pseudomonas* and *Burkholderia* likely contributed to the rapid removal of less recalcitrant compounds, caffeine, and paracetamol, within seven days (Fig. 3). Both genera are known to harbor oxygenases and demethylases involved in paracetamol and caffeine metabolism (Lin et al., 2023; Wu et al., 2012), even though these genes were not captured in the MAGs. The genus *Achromobacter*, abundant during initial cultivation and carrying *linE*, was also likely involved in paracetamol degradation (Fig. 3).

Metagenomic analysis of MBR1 and MBR6 revealed a repertoire enriched in monooxygenases and dioxygenases (Fig. 5, Supplementary Materials S4), supporting the predominance of oxidative degradation mechanisms in pharmaceutical removal. These enzyme classes are known for their broad substrate range and catalytic versatility, confirming the biochemical potential of adapted communities for micropollutant transformation (Stadlmair et al., 2018).

However, the interpretation of gene abundance data remains limited. Quantitative differences in gene counts did not consistently mirror reactor performance, emphasizing that gene presence alone does not capture true degradation potential. This limitation largely arises from the incomplete understanding of many pharmaceutical degradation pathways, which constrains accurate annotation of the responsible genes (Borchert et al., 2021).

The progressive increase in community diversity over time implies that broader microbial diversity can enhance degradation efficiency, an outcome opposite to what was initially hypothesized. This may result from more extensive and promiscuous enzymatic machinery capable of acting on multiple classes of pollutants (Hernandez-Raquet et al., 2013).

Despite the positive removal results, the system exhibited clear limitations. Diclofenac was substantially more recalcitrant to biodegradation in the lab-scale MBR than the other substrates, likely because its chlorinated structure hinders the hydroxylation step required to initiate degradation(Bouju et al., 2016). This challenge aligns with the limited number of studies reporting efficient diclofenac biodegradation (Demaria et al., 2026; Ivshina et al., 2019; Yang et al., 2022): one study reported that only 30% was removed under anaerobic conditions over 24 days (Yang et al., 2022), and another one that 60% was removed over 20 weeks in an MBR inoculated with a thermophilic compost-derived consortium (Demaria et al., 2026). In contrast to the lab-scale performance, the 7-L MBR treating biologically treated wastewater showed a clear decrease in diclofenac concentrations after 32 days of operation, indicating that scaled-up conditions enabled partial degradation of this recalcitrant compound. The scaled-up system achieved near-complete removal of caffeine, enalapril, ibuprofen, and paracetamol, while atenolol and diclofenac were removed by up to 75% and 50%, respectively, within 31 days (Fig. 6a). When the adapted microbial community from MBR6 was introduced into the 7-L reactor containing pharmaceuticals at ng-µg/L in a 7 L reactor, it also exhibited functional adaptation to compounds absent during the laboratory enrichment phase, demonstrating its flexibility and metabolic versatility. The community degraded several untargeted compounds detected in the effluent, including naproxen, ketoprofen, valsartan, and nicotine, achieving 75–100% removal (Fig. 6b). The target pharmaceuticals investigated in this study, atenolol, caffeine, paracetamol, ibuprofen, and enalapril, exhibit strong structural similarities to several non-target compounds, which likely facilitated their concurrent transformation by the adapted microbial consortia. Most of these molecules contain an aromatic ring (atenolol, enalapril, paracetamol, ketoprofen, naproxen, valsartan). A benzene ring can be microbially activated through hydroxylation to intermediates that subsequently undergo intra-, or estradiol-cleavage catalyzed by dioxygenases (Fuchs et al., 2011). In addition, many pharmaceuticals include carbonyl-containing groups (such as amides, esters, or carboxylic acids) that serve as reactive electrophilic centers, enabling hydrolysis and oxidation to initiate degradation (Dey, 2015). Compounds like caffeine, nicotine, and enalapril while structurally distinct, possess nitrogen heterocycles (pyrrolidine, pyridine, or imidazole-like rings) that are electron-rich and readily oxidized, allowing comparable ring activation and opening mechanisms (Schwarz and Lingens, 1994). Thus, the shared presence of aromatic systems, carbonyl groups, and oxidizable heteroatoms could explain the degradation of both target and non-target pharmaceuticals through similar enzymatic activation and cleavage pathways.

In summary, the adapted microbial community retained its performance in real wastewater, efficiently removing both targeted and previously unexposed pharmaceuticals. These findings demonstrate the ecological resilience, functional versatility, and scalability of micropollutant-adapted microbiomes as biological polishing stages in tertiary treatment, aimed at mitigating pharmaceutical release in the environment.

## ACKNOWLEDGMENT

This work was supported by the European Union’s Horizon Europe program (Grant 101060625 – NYMPHE) and the Swiss State Secretariat for Education, Research and Innovation (SERI). Additional support was provided by the “César Nombela” Research Talent Attraction Program (Grant 2024-T1/ECO-31227, Comunidad de Madrid, R.B.).

## ABBREVIATIONS

WWTP: wastewater treatment plant
MBR: membrane bioreactor
HRT: hydraulic retention time
COD: chemical oxygen demand
HPLC: high performance liquid chromatography
ASV: amplicon sequence variant
NMDS: non-metric multidimensional scaling
DESeq: Differential abundance analysis
MAG: Metagenome-assembled genomes
LC-MS: liquid chromatography-mass spectrometry
MS: mass spectrometry
ESI: electrospray ionization
MRM: multiple reaction monitoring

## Notes

### Competing Interest Statement

The authors have declared no competing interest.

